# Male Mice Have no Sugar-preference Behavior

**DOI:** 10.1101/2024.12.11.627417

**Authors:** Tao Pan, Jia-Fu Huang, Qi-Ci Wu, Yi-Xuan Chen, Xiu-Min Li, San-Guo Zhang, Yu-Tian Pan

## Abstract

Sugar-preference behavior is considered to be universally innate among humans and animals. Current research on the detection and mechanisms of sugar-preference behavior largely adopts the “two-bottle choice assay”, and most experiments use male mice as the model, which has serious methodological flaws, leading to questionable conclusions. As expected, through adopting our improved experimental method, we discovered distinct sugar preferences between male and female mice: females possess an innate preference, while males do not. However, the innate sugar preference in female mice can be lost due to their long-term normal diet. Even after “training” with high-sugar food for a long time, the results still show differences between males and females. Female mice retain a significant preference for sugar, while male mice are not influenced by the training of high-sugar food and still do not evolve a sugar preference. From this, we conclude that gender determines the emergence and development of sugar preference in mice. Our results can provide researchers with a new understanding of sugar preference, which will help them fill in or even correct the gaps in the original experimental design methods to obtain more reliable conclusions.

---

Sugar is a vital component of the human diet, but excessive consumption has long been associated with numerous health risks, mainly including obesity, diabetes, cardiovascular disease, osteoporosis, inflammatory bowel disease, neurological disorders, and certain cancers ^[1-6]^. In response, the World Health Organization (WHO) has issued guidelines recommending a daily sugar intake of no more than 25 grams to mitigate these risks ^[6]^. Despite this, a strong preference for sugar is pervasive among both humans and animals, prompting significant scientific interest in understanding its underlying mechanisms.

Nelson *et al*. ^[7]^ demonstrated that non-thirsty wild-type mice offered a choice between water and a sugar solution predominantly consumed the sugar solution within 48 h. Notably, animals can exhibit a strong preference for sugar even in the absence of functional sweet taste receptors ^[8-10]^, suggesting that there is a mechanism for sugar preference independent of taste. As sugar serves as a fundamental energy source for animals, most species have evolved specialized brain circuits that facilitate the seeking, recognition, and stimulation of sugar consumption ^[11]^. Building on this, Tan *et al*. ^[12]^ identified the gut-brain axis as a key pathway mediating sugar-preference behavior. These findings appear to provide insight into the biological basis for sugar preference in animals. However, in a recent study, we conducted a simple animal behavior experiment (Fig. 1A, and Fig. 2) that yielded unexpected results.

**Fig. 1.**
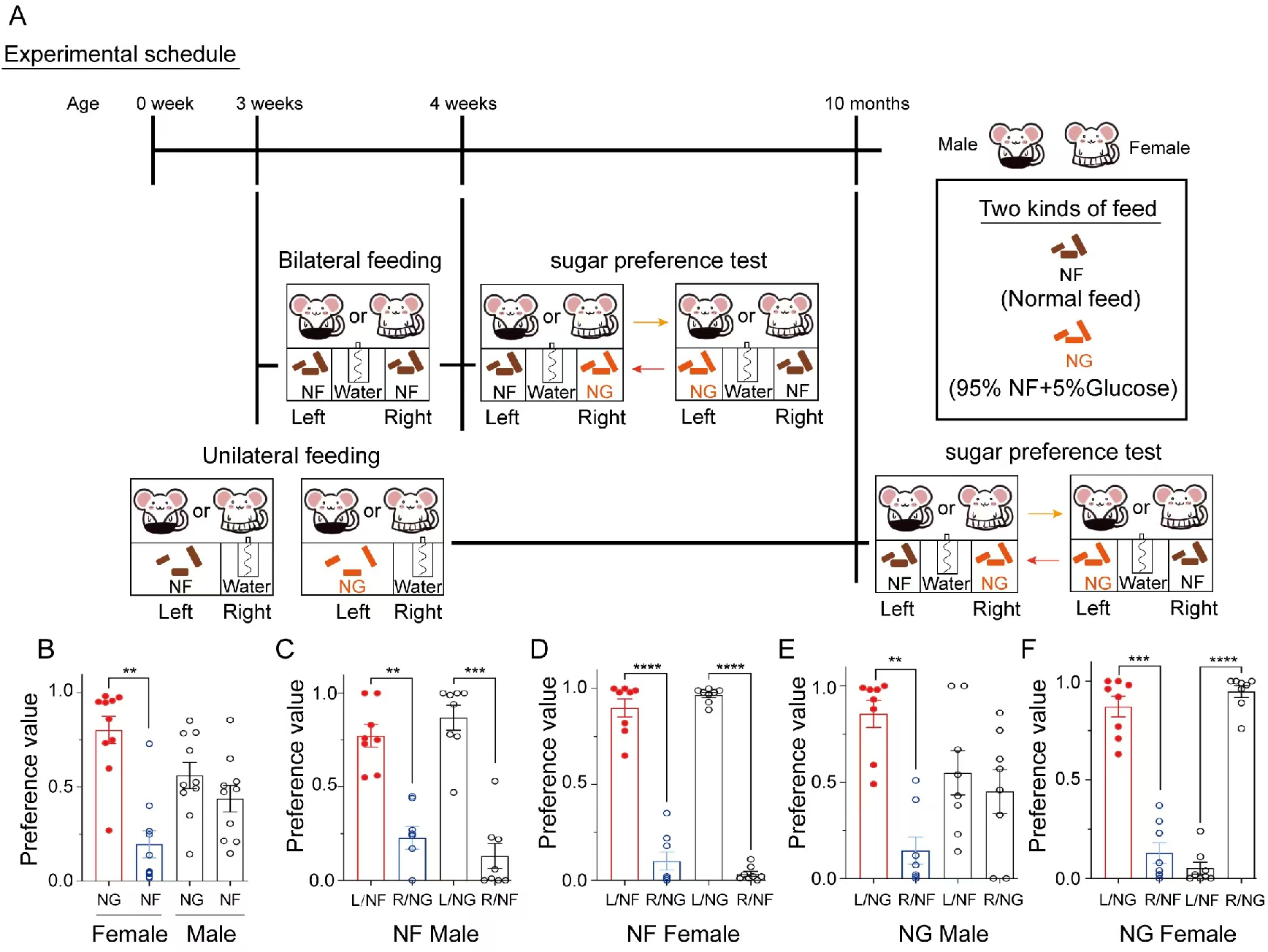
Sugar preferences in male and female mice. (**A**) Experimental timeline for sugar-preference testing. (**B**) Feeding preference of young male and female mice for NF versus NG. Female mice displayed a significant sugar preference, whereas male mice showed no significant difference. (**C** to **F**) Feeding preference of mice conditioned to consume a single feed (NF or NG) placed on the left side for an extended period, tested with both NF and NG provided simultaneously.

**Fig. 2.**
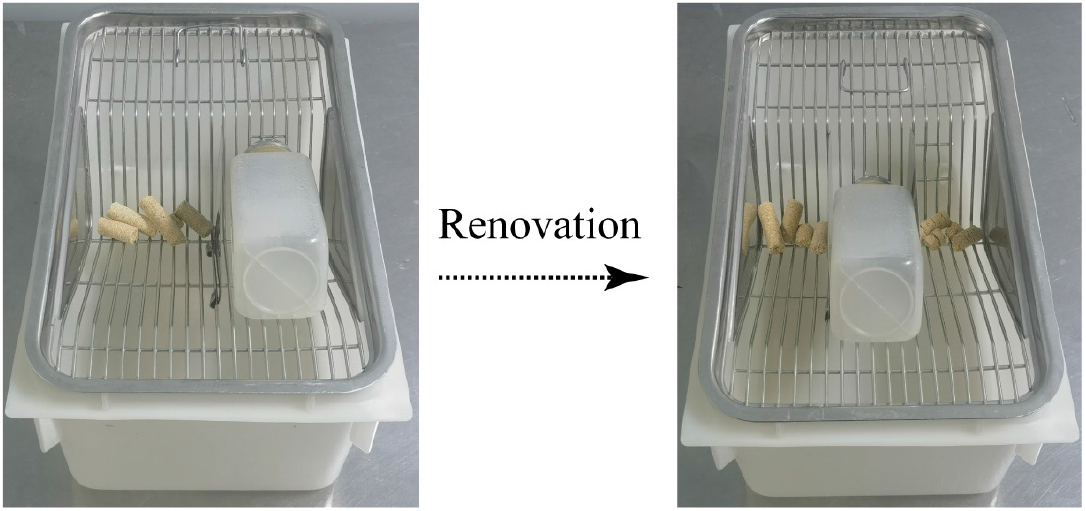
Modifications to the feeding cage design. (**A**) Standard cage setup, with feed placed on the left and the water bottle on the right. (**B**) Modified cage setup, with the water bottle centrally located and feed placed on both the left and right sides to minimize positional bias during preference testing.

We first examined sugar-preference behavior in young mice. Over a 48-h period, male and female mice with no prior exposure to high-sugar diets exhibited distinct feeding preferences (Fig. 1B). Female mice displayed a significant preference for high-sugar feed (NG) over normal feed (NF) (n=10, paired *t*-test, *P*_Female_=0.0023). In contrast, male mice showed no significant preference differences in their consumption of NF and NG (n=10, paired *t*-test, *P*_Male_=0.40). These findings were consistent across 24-h intervals, as confirmed by independent statistical analyses (Fig. 3). Therefore, young male mice do not demonstrate innate sugar-preference behavior, unlike their female counterparts.

**Fig. 3.**
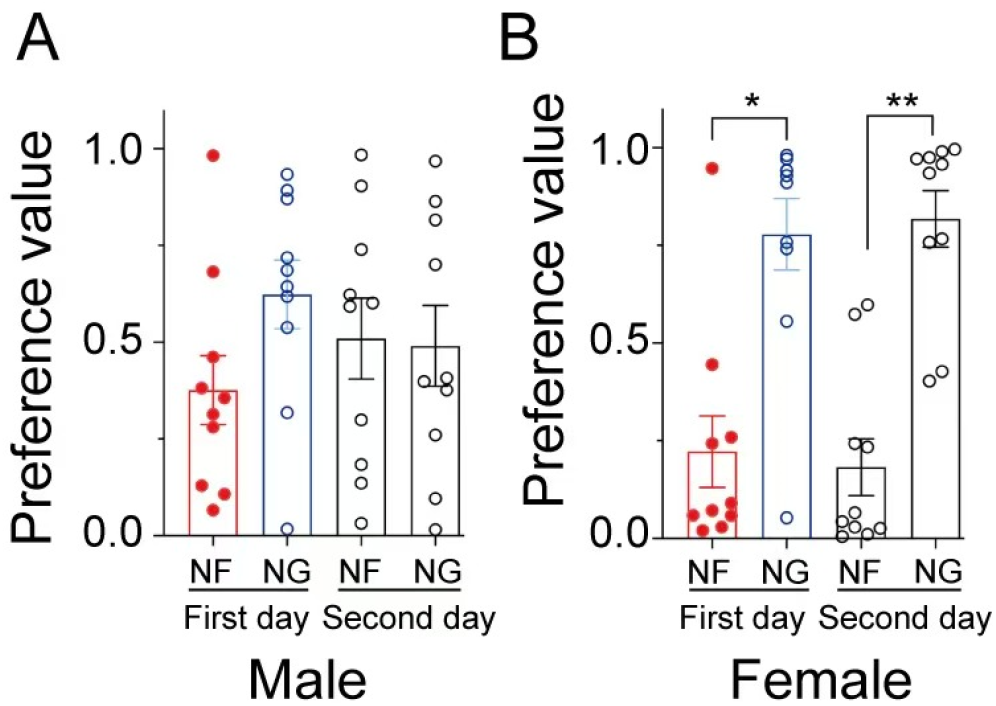
Initial sugar-preference testing in young mice. (**A**) Initial sugar-preference tests in young male mice. No significant sugar preference was observed on day 1 or day 2 (n=10, *P*_*male*_=0.20 and *P*_*male*_=0.93, respectively). (**B**) Initial sugar-preference tests in young female mice. Significant sugar preference was detected on both two days (n=10, *P*_*female*_=0.014 on day 1 and *P*_*female*_=0.0018 on day 2).

Next, we investigated how dietary habits influence sugar-preference behavior. Mice underwent long-term (10-month) training with unilateral (left-side) feeding of either NF or NG, resulting in clear habitual preferences (Fig. 4). During sugar-preference testing, distinguishing between habitual preferences and innate sugar preference was essential to avoid misinterpretation. When NG was placed in the habitual feeding position (left) and NF in the non-habitual position (right), all mice strongly preferred NG (Fig. 1, C to F; *P*_*NF male*_= 0.0009, *P*_*NF female*_ < 0.0001, *P*_*NG male*_= 0.0015, *P*_*NG female*_ < 0.0001). When the NF was positioned in the habitual feeding position (left) and NG in the non-habitual position (right), behavior varied by gender and diet group. In the NF-trained group, both male and female mice strongly preferred NF at the habitual position (Fig. 1, C and D; n=8, paired *t*-test, *P*_*NF male*_=0.0027. *P*_*NF female*_ < 0.0001). However, in the NG-trained group, male mice showed no significant preference between NF and NG, regardless of position (Fig. 1E; n=8, paired *t*-test, *P*_*NG male*_=0.68). Female mice in the NG-trained group, however, demonstrated a strong preference for NG at the non-habitual position (right) over NF at the habitual position (left), prioritizing sugar preference over feeding habit (Fig. 1F; n=8, paired *t*-test, *P*_*NG female*_ < 0.0001). These results suggest that male mice on a normal diet do not exhibit sugar-preference behavior, instead favoring the habitual feeding position. Even after long-term high-sugar diet training, male mice failed to develop a preference for sugar. In contrast, female mice displayed distinct behavioral patterns. Female mice on a daily normal diet lost their innate sugar-preference behavior and developed a clear preference for the habitual feeding position. However, females subjected to prolonged high-sugar diet training retained a strong sugar-preference behavior, overriding habitual feeding tendencies.

**Fig. 4.**
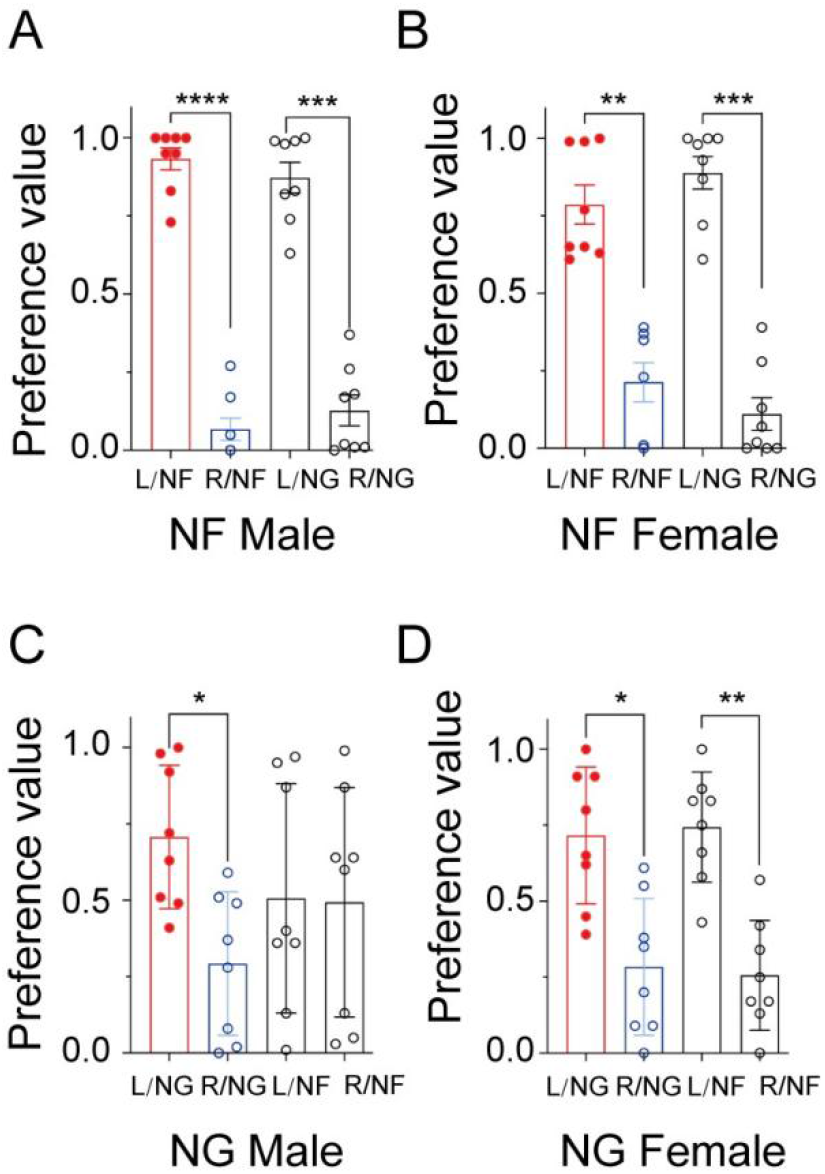
Influence of feed type and position on feeding habit preference. (**A**) Male mice fed NF exclusively on the left side over a prolonged period showed a strong positional preference for the left side, consuming either NF or NG (n=8, *P*_*NF*_<0.0001; *P*_*NG*_=0.0001). (**B**) Female mice fed NF exclusively on the left side also displayed a significant positional preference for the left side with both NF and NG (n=8, *P*_*NF*_=0.0026; *P*_*NG*_=0.0001). (**C**) Male mice conditioned on NG placed on the left side demonstrated a positional preference for NG on the left (n=8, *P*_*NG*_=0.041). No significant preference was observed for NF consumption between the left and right sides (n = 8, *P*_*NF*_ = 0.96). (**D**) Female mice conditioned on NG on the left side exhibited a positional preference for both NG and NF on the left (n=8, *P*_*NG*_=0.030; *P*_*NF*_=0.0066).

Our experimental findings provide new insights for researchers studying sugar preference, addressing gaps in current experimental design and offering more reliable conclusions. For example, in the study by Nelson *et al*. ^[7]^, a “two-bottle choice assay” was employed to assess sugar preference. In this assay, 2–3 mice were housed together and given simultaneous access to two bottles—one containing a sugar solution and the other water. Sugar preference was inferred if the sugar solution accounted for more than 50% of total intake over 48 h. However, this setup inherently limits free-choice behavior, as only one drinking spout per bottle is available, conditioning individual drinking choices. Moreover, the group setting precludes accurate tracking of individual consumption. Even if total sugar solution intake exceeded water intake, the method cannot confirm that all mice in the cage exhibited sugar-preference behavior. Unfortunately, the findings of this study have been widely accepted, forming the basis for subsequent research into sugar preference mechanisms. As a result, studies exploring the mechanisms of sugar preference may have relied on an uncertain or flawed model, undermining the reliability of their conclusions. Additionally, a review of existing literature reveals a striking oversight: most studies omit data on female mice, often generalizing findings from male mice to all “mice.” This study highlights an important distinction: sugar preference differs between sexes. Using male mice as the sole model for sugar-preference research risks significant errors or omissions. Female mice maintained on high-sugar diets provide a more appropriate model for investigating the underlying mechanisms of sugar preference.

In summary, innate sugar preference in mice is gender-specific: females exhibit this behavior while males do not. However, prolonged exposure to a normal diet can extinguish the innate sugar preference in female mice. Furthermore, the impact of long-term exposure to high-sugar diets differs by gender. Female mice retain a strong preference for sugar, whereas male mice remain unaffected and fail to develop such a preference. Thus, gender plays a critical role in the development and persistence of sugar preference in mice.

These findings have significant implications for dietary health. To mitigate the risks associated with high-sugar diets, altering the location of daily eating habits or modifying the placement of high-sugar foods may help reduce the intake of sugars in “gentlemen”. However, for “ladies”, effective risk management may require removing high-sugar foods from their environment as much as possible.

## Materials and methods

### Laboratory mouse

Male and female ICR mice (SPF grade) were housed under specific pathogen-free conditions in a temperature-controlled room (25 °C) with 50-60% humidity and a 12-h light/dark cycle. Autoclaved water and food were provided *ad libitum*. All experimental procedures were reviewed and approved by the Animal Ethics and Welfare Committee of Minnan Normal University (approval No. AEWC- 2023010, Zhangzhou, China).

### Feeding cage modification

Standard mouse feeding cage, which typically position feed on the left side and a water bottle on the right side, were modified for this study. The water bottle was relocated to the center, and experimental feeds were positioned on both the left and right sides. This modification was implemented to standardize adaptive feeding behavior prior to preference testing and to eliminate positional bias in feed preference assessments (Fig. 2).

### High glucose feed preparation

Based on findings by Nelson *et al*. ^[7]^, who reported significant sugar-preference behavior in mice at sugar concentration of 40-320 mM, a median concentration of 5 % sugar was selected for this study. In fact, glucose is also commonly used in sugar-preference studies ^[12]^. Thus, high-glucose feed (NG) was prepared by supplementing normal feed (NF) with 5 % glucose, following the feed preparation protocol outlined by Shinobu Hirai.*et al* ^[13]^.

### Behavioral testing

#### Detection of sugar-preference behavior

The preference behaviors were assessed by individually housing each mouse in a modified cage. During the first 24 h, NF was placed on one side and NG on the opposite side. Feed positions were then swapped, and feeding behaviors were observed for an additional 24 h. The intake from each side was weighed and recorded daily to quantify feeding preferences.

#### Detection of sugar-preference behavior in young mice

Three-week-old weaned male and female mice were selected and trained to consume the same feed (NF) placed on both sides of the cage for 7-10 days before testing. This training was conducted to eliminate the potential of the preference of habitual feeding position. The sugar-preference behavior was then assessed using the methodology as described above.

#### The effects of dietary conditions on the development of sugar-preference behavior in mice

Three-week-old weaned mice, equally divided by gender, were randomly assigned to two dietary groups. Mice in the NF group were fed NF placed on left side of the cage, while those in the NG group were fed NG placed on left side. After 10 months of feeding, the habitual feeding position preference and sugar-preference behavior were assessed as the aforementioned methods.

#### Detection of the preference for feeding position

To evaluate feeding position preferences, the same kind of feed (NF or NG) was placed on both sides of the cage. Feed intake was observed and recorded for 24 h. The feed was then replaced with the alternate type, and feeding behaviors were observed for another 24 h. The intake from both the left and right sides was weighed and recorded daily.

#### Statistical analysis

The above bilateral preference values were analyzed using *t*-test. Data are expressed as mean ± S.E.M. The preference value was calculated as follows: Feeding preference value for the habitual position = Feed intake on the left side / (Feed intake on the left side + Feed intake on the right side); Sugar-preference value = Feed intake of high-sugar feed / (Feed intake on the left side + Feed intake on the right side).

## Acknowledgments

We are grateful to the members of Pan Laboratory from Minnan Normal University for the discussion and technical assistance. This work was financially supported by the National Science and Technology Planning Project of China (No. 2021L3027), the General Program of the National Natural Science Foundation of China (No.32370888).

